# The conserved coordination of acyl-homoserine lactone and PqsE signaling defines the RhlR-dependent quorum-sensing network in *Pseudomonas aeruginosa* clinical isolates

**DOI:** 10.64898/2026.05.21.726854

**Authors:** Elizabeth G. Knorr, Elizabeth A. Key, Megan L. Schumacher, Leah M. Kemper, Caleb P. Mallery, Navjot Singh, Jon E. Paczkowski

**Affiliations:** Division of Genetics, Wadsworth Center, New York State Department of Health, Albany, New York, USA; Department of Biomedical Sciences, University at Albany, College of Integrated Health Sciences, Albany, New York, USA; Division of Scientific Cores, Wadsworth Center, New York State Department of Health, Albany, New York, USA; The RNA Institute, University at Albany, College of Arts and Sciences, Albany, New York, USA

## Abstract

Quorum sensing (QS) enables *Pseudomonas aeruginosa* to coordinate virulence and biofilm formation through cell density-dependent signaling. In clinical isolates from patients with cystic fibrosis (pwCF), mutations in canonical QS systems such as *lasR* and *rhlI* often lead to altered signaling hierarchies that complicate our understanding of QS regulation during chronic infection. Here, we dissect the relative contributions of the autoinducer *N*-butyryl-L-homoserine lactone (C_4_HSL) and the protein binding partner PqsE to RhlR-dependent transcription in CF clinical isolates. Using site-directed mutagenesis to generate RhlR and PqsE variants incapable of responding to C_4_HSL (RhlR A44M) or dimerizing to interact with RhlR (PqsE^NI^), we show that both inputs are essential for the full expression of QS-regulated virulence factors, including pyocyanin and rhamnolipids. Transcriptomic analyses revealed that C_4_HSL and PqsE co-regulate a conserved set of 28 RhlR-dependent genes, encompassing canonical virulence loci as well as uncharacterized genes that are likely important for adaptation to the CF airway environment. These findings establish that clinical isolates maintain functional QS circuitry reliant on dual activation of RhlR by both C_4_HSL and PqsE, revealing a conserved regulatory module that underpins pathogenic behavior across genetically diverse isolates.

**AUTHOR SUMMARY:** Understanding quorum-sensing regulation in clinical isolates of *Pseudomonas aeruginosa* is essential to determine how the pathogen persists and adapts within the cystic fibrosis lung. While most studies have focused on laboratory strains, chronic isolates exhibit distinct genetic and regulatory adaptations that complicate our ability to generalize quorum sensing function. Our work defines the coordinated roles of C_4_HSL and PqsE in activating RhlR-dependent gene expression and virulence factor production in isolates from patients with cystic fibrosis. We identify a conserved core of quorum-sensing-regulated genes that remain dependent on both signals despite extensive genomic divergence. These findings highlight that, even within the evolutionary landscape of chronic infection, quorum-sensing signaling through RhlR remains a central and conserved determinant of virulence. By resolving the dual contributions of acyl-homoserine lactone and PqsE-mediated activation, this work provides a mechanistic foundation for future efforts to therapeutically target quorum-sensing pathways in clinical *P. aeruginosa* infections.

## INTRODUCTION

Cystic fibrosis (CF) is a genetic disorder caused by mutations in the gene encoding the CF transmembrane conductance regulator (CFTR) protein [1–3]. CFTR variants cause a thickening of the mucus in the airways and impaired mucociliary clearance leading to underlying respiratory dysfunction and increased vulnerability to infection [4–7]. *Pseudomonas aeruginosa* is an opportunistic pathogen that commonly infects the airways of patients with CF causing significant morbidity and mortality [8–12]. Patients with CF (pwCF) are often infected with *P. aeruginosa* early in life with subsequent re-infections before developing a chronic infection, which can no longer be eliminated by traditional antibiotics [10, 13–19]. These chronic infections can evade host immune defenses and antibiotic treatment through acquired mechanisms of multidrug resistance and intrinsic mechanisms of resistance such as biofilm development [17, 20–25].

Quorum sensing (QS) is a density-dependent cell-to-cell communication mechanism employed by *P. aeruginosa* to detect and respond to the presence of additional *P. aeruginosa* via secreted small molecules called autoinducers (AI), leading to cooperative group behaviors [26–33]. An autoinducer binds to its respective receptor to facilitate QS via alterations in gene expression, which regulate biofilm formation and development, virulence factor production, and pathogenesis [33–37]. QS in *P. aeruginosa* is primarily driven by the LuxI/R synthase/receptor pairs, LasI/LasR and RhlI/RhlR. LuxR-type receptors possess a variable N-terminal ligand binding domain that contains a bipartite ligand binding pocket (LBP) that can accommodate acyl-homoserine lactones and a C-terminal helix-turn-helix DNA-binding domain [38]. LasI synthesizes 3OC_12_HSL which binds to LasR leading to the transcription of QS genes including *rhlI* and *rhlR* [28, 30, 34, 39–42]. RhlI synthesizes C_4_HSL, which binds to the receptor RhlR, inducing the expression of additional QS genes [26, 30]. This positive feedback loop in response to AI levels enables *P. aeruginosa* to coordinate gene expression that underpins virulence factor production, biofilm formation, and pathogenesis, among other group behavior traits [34, 43, 44]. We recently discovered a RhlR variant (A44M) via site-directed mutagenesis of residues in the LBP that was incapable of being activated by C_4_HSL but was still soluble when expressed in an *Escherichia coli* heterologous system or natively from the *rhlR* locus in *P. aeruginosa* [45], representing a unique genetic tool to directly assess RhlR-ligand-dependent activation *in vivo* and in clinical isolates.

In addition to the Las/Rhl QS pathways, the Pqs system employs a similar feedback mechanism with the receptor, PqsR, binding the AI PQS to upregulate the *pqsABCDE* operon, which encodes for the enzymes in the PQS biosynthetic pathway [46–53]. It was recently discovered that PqsE, the final product of the *pqs* operon, is required for virulence factor production and pathogenesis [54–60]. However, we showed that this dependency is not through its role as an enzyme in the PQS biosynthetic pathway; rather, it is through a physical interaction with RhlR [61–65]. RhlR bound by PqsE exhibited enhanced stability and DNA binding capability [61, 62]. The influence of PqsE on RhlR-dependent DNA binding and gene expression is associated with its ability to dimerize [62, 66]. Indeed, we recently showed that *P. aeruginosa* PqsE dimerization is unique to this class of proteins [66]. Disruption of the PqsE dimerization interface via three arginine residues substituted to alanine, which we term the PqsE^NI^ (non-interacting) variant, resulted in an inability to properly regulate RhlR function [62, 66]. PqsE protein levels are tightly controlled and correlate inversely with C_4_HSL-dependent activation of RhlR, as activation of RhlR, either by high levels of AI or *rhlR* overexpression, represses *pqsABCDE* gene operon expression [45, 67].

Clinical isolates of *P. aeruginosa* from both chronic and acute infections are well-known to carry mutations that result in altered QS signaling [68–72]. We recently isolated common mutations from these strains and cloned them into their native locus in the laboratory strain UCBPP-PA14 (herein referred to as PA14). We found that laboratory strains of *P. aeruginosa* harboring *lasR* mutations rendered the *las* system non-functional but produced nearly triple the amount of C_4_HSL compared to the wild-type (WT) strain [73]. Counterintuitively, pyocyanin production, an important toxin that is controlled by QS via RhlR in these strains, was repressed, indicating that high levels of C_4_HSL can alter RhlR activity [73, 74]. RhlR signaling was restored when common, co-occurring *rhlI* mutations were introduced into the *lasR* mutant backgrounds due to the re-calibration of C_4_HSL levels to WT concentrations [73]. Thus, C_4_HSL levels play an important role in mediating QS progression and there is a multi-step process to activating RhlR that has not been fully characterized in either laboratory strains or clinical isolates. Previous work exploring the role of RhlR on QS in laboratory strains and clinical isolates used full gene deletions, which obscures the contributions of individual co-regulators of RhlR activity [57, 62, 75–80]. Here, we specifically assess the individual contributions of C_4_HSL and PqsE to QS signaling and virulence factor production by introducing into clinical isolates a *rhlR* mutation that results in a protein variant incapable of responding to C_4_HSL (*i.e.*, A44M) or a *pqsE* mutation that results in a variant incapable of binding to and regulating RhlR (*i.e.*, PqsE^NI^) [45, 62]. We found that disruption of the ability of RhlR to bind C_4_HSL or PqsE resulted in significant alterations in QS gene expression. We define the contributions of both C_4_HSL and PqsE to RhlR-dependent gene expression, establishing a core regulon among laboratory strains and clinical isolates. Furthermore, we establish the individual and shared roles of C_4_HSL and PqsE in contributing to RhlR-dependent pathogenesis of clinical isolates in a mammalian cell culture infection model.

## RESULTS

### Whole genome sequencing of isolates reveals divergent isolates with intact QS networks and key polymorphisms

We performed a comparative genomic analysis of six clinical *P. aeruginosa* isolates obtained from pwCF (JPS1005, JPS1006, JPS1008, JPS1009, JPS1010, JPS1011) against the UCBPP-PA14 (PA14) reference genome to identify regions of chromosomal divergence and to detect isolate-specific elements (**Figure1, Table S1**). All isolates were within the expected *P. aeruginosa* genome size range and GC content (approximately 6.3–7.1 Mb with approximately 66% GC) (**Table S1**). JPS1005 and JPS1006 had the largest genomes relative to PA14 with an approximate acquisition of 315,000 and 560,000 bp, respectively,. Meanwhile JPS1008, JPS1009, and JPS1010 displayed modest genome reduction with an average difference of approximately 142,500 bp, likely consistent with the loss of PA14-specific regions. Comparatively, JPS1011 differed in total genome size from PA14 by only ∼20,000 bp (**Table S1**). Despite variation in total size, all isolates had a conserved core PA14-like genome. We mapped regions of low nucleotide similarity between PA14 and each isolate using a 10-kb sliding window (**Table S2**). Overall, there were 17 regions of greater than 3,000 bp that were missing in all isolates compared to PA14. In particular, we found two major regions of divergence: (i) a prophage island near 1.92-1.94 Mb encoding for canonical phage structural genes with low GC content for *Pseudomonads* (approximately 51%) and (ii) an O-antigen/LPS biosynthetic locus near 2.03-2.05 Mb encoding glycosyltransferases, flippases, and O-antigen polymerases; (**Figure 1**). These regions likely represent genomic islands characterized by mobility genes and functions subject to environmental and host-driven selection. Gene-level presence and absence matrices revealed that none of the coding sequences in the PA14 islands were present as exact nucleotide matches in any clinical isolate, indicating that the isolates lack the PA14 prophage and O-antigen cluster. Their absence likely reflects replacement, deletion, or substantial sequence remodeling rather than complete loss of biological functions, as surface modification, mobile element activity, and horizontal gene transfer are known to be variable among *P. aeruginosa* lineages.

**Figure 1.**
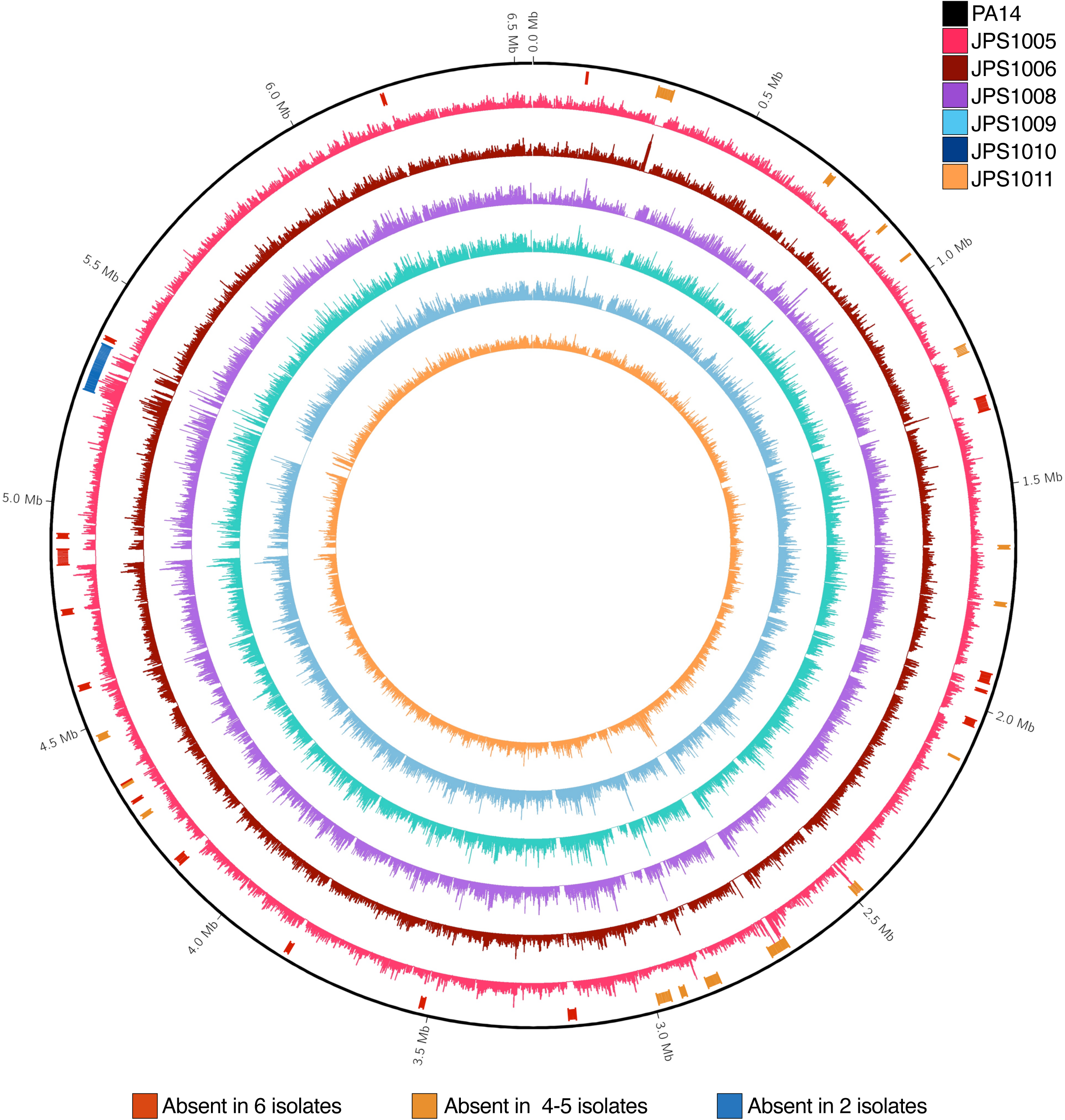
Whole genome sequencing of isolates reveals divergent isolates with intact QS networks and key polymorphisms. Circos plot displaying the distribution of SNPs across the genome of each isolate aligned to the UCBPP-PA14 reference genome (black). The plot was constructed using the results of variant calling from whole genome sequencing of each isolate. The number of SNPs was counted with bin size of 2000 bp to create a circular histogram plot. Each track represents the genome of the designated isolates.

To identify isolate-specific accessory regions, we reversed the comparison by scanning each clinical genome for windows with minimal similarity to PA14. This revealed distinct sets of horizontally acquired elements in each isolate. These windows exhibited altered GC content, consistent with recent acquisition via phage or integrative conjugative elements. For example, JPS1011 showed divergence near 4.91-4.93 Mb (**Table S3**), supporting independent acquisition of mobile elements within the clinical environment. Overall, these results demonstrate that the major drivers of genomic divergence between PA14 and this cohort of clinical isolates are in accessory genome remodeling. The core chromosome is highly conserved, while accessory regions exhibited lineage-specific gain, loss, and replacement. These results highlight the dynamic nature of *P. aeruginosa* genome evolution in clinical contexts and provide a foundation for downstream functional and phenotypic correlation of QS traits.

Due to the frequency of mutations in the QS network of isolates obtained from pwCF, we investigated the sequences of key QS regulators and their downstream targets that mediate community-wide behaviors. We found there was genetic diversity in the QS networks among the isolates obtained from pwCF (**Figure 1**). Importantly, for this study, only synonymous mutations were found in *rhlR* and *pqsE* (**Table 1**). We identified several missense, insertions, deletions, and frameshifts in other QS components, including *rhlI*, *pqsR* (*mvfR*), and *pqsABD* and *pqsH* (**Table 1**, **Figure S1**). Additionally, we identified many SNPs in QS outputs regulated by the RhlR-PqsE system, including *lasA*, *lasB*, *rhlA*, *rhlB*, *mex*, *lecB*, *phnA*, and *hcnA* (**Table 1**, **Figure S1**). Lastly, motility genes (genes encoding proteins involved in: pilus formation such as *pilB*, flagella formation such as *flgE*, and chemotaxis such as *cheA*) and biofilm genes (genes encoding cyclic-di-GMP synthases such as *wsp* and genes encoding matrix components such as *pel*, *psl, muc,* and *alg*) were highly enriched for SNPs (**Table 1, Figure S1**). Given the conserved sequences of *rhlR* and *pqsE* and the important role of the RhlR-PqsE module in driving infections, we sought to determine their role in regulating QS in the context of divergent clinical isolates.

**Table 1.** SNPs of QS genes.

### The PqsE-RhlR interaction regulates pyocyanin production in isolates from pwCF

To determine the role of PqsE-dependent activation of RhlR in isolates obtained from pwCF, we engineered mutations at the native locus for *pqsE*. The amino acid sequence of PqsE for each of the isolates was identical to that of PA14. To test that PqsE dimerization and subsequent interaction of PqsE dimers with RhlR was maintained in these isolates, we measured pyocyanin production in our cohort of clinical isolates expressing PqsE^NI^, which is a PqsE variant incapable of dimerizing and forming a complex with RhlR. The parent strains obtained from pwCF produced variable amounts of pyocyanin (**Figure 2A**). JPS1004, JPS1008, and JPS1009 produced pyocyanin comparable to that of PA14, while JPS1010 and JPS1011 produced 2-fold higher levels of pyocyanin compared to PA14. Strain 1006 was the outlier in our cohort, as it produced relatively little pyocyanin. We hypothesize that this is due to mutations in genes required for pyocyanin production, namely *phzS, phzH*, and *phzM* (**Figure S1**, **Table 1**). Importantly, disruption of the PqsE dimerization interface and, thus, its ability to interact with RhlR to upregulate transcription at the phenazine promoters, abolished the ability of all isolates from pwCF to produce pyocyanin (**Figure 2A**). PqsE and C_4_HSL exert similar effects on RhlR-dependent pyocyanin production, and the presence of both are necessary for pyocyanin production. To confirm that the decrease in pyocyanin production was due to the inability of PqsE to interact with RhlR and not due to a decrease in C_4_HSL levels, we measured the levels of AI produced by the parental clinical isolates and their corresponding *pqsE^NI^* mutants using mass spectrometry (**Figure 2B**). C_4_HSL concentrations remained unchanged between the parent strain and their respective partner strains expressing the PqsE variant (**Figure 2B**). Interestingly, JPS1004 exhibited elevated C_4_HSL production relative to WT PA14 (**Figure 2B**), but this did not result in elevated levels of pyocyanin production by this strain (**Figure 2A**), indicating that clinical isolates from pwCF exhibit control of QS traits to prevent excess virulence factor production, similar to what has been observed in laboratory strains [45, 55, 57, 73]. Collectively, these data indicate that specifically disrupting the RhlR-PqsE interaction in clinical isolates leads to a decrease in pyocyanin production.

**Figure 2.**
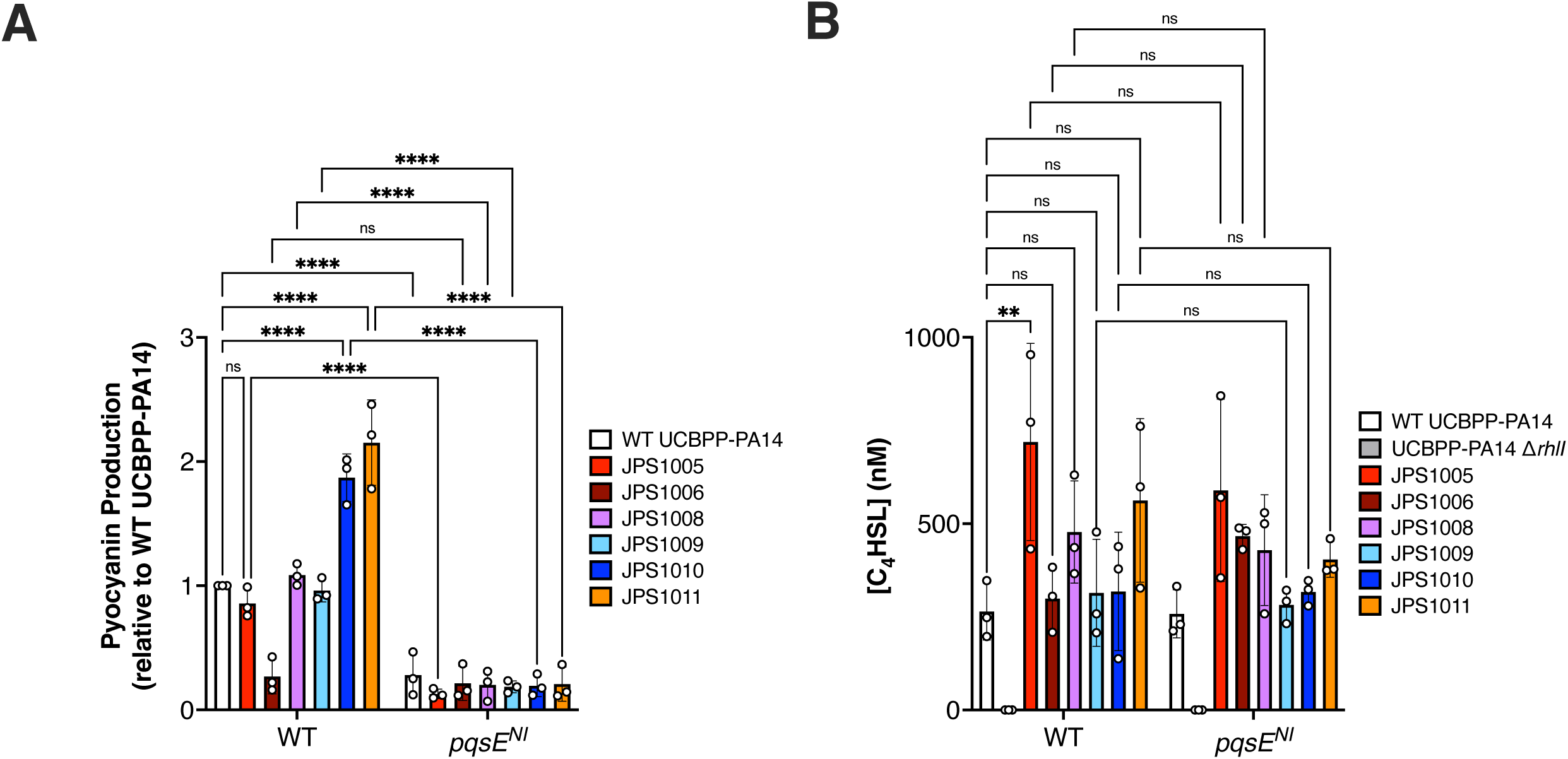
PqsE is essential for pyocyanin production in clinical isolates. **A.** Pyocyanin production as measured by OD_695 nm_ / OD_600 nm_ for WT UCBPP-PA14 (white) and different clinical isolates as well as their isogenic *pqsE^NI^* variants. All data were normalized to the pyocyanin produced by UCBPP-PA14 WT. **B.** Absolute concentrations of C_4_HSL from cell-free supernatants of the same strains shown in (**A**) as well as a 11*rhlI* (gray; C_4_HSL negative control) as measured by UHPLC-HRMS. Statistics were performed using a 2-way analysis of variance (ANOVA) with a Šidák’s multiple comparisons test. p-value summary: **** < 0.0001; ns = not significant.

### RhlR sensitivity to C_4_HSL controls pyocyanin production in isolates from pwCF

Based on our current understanding of *P. aeruginosa* QS, activation of RhlR by PqsE is only one-half of the inputs that control RhlR-dependent expression of phenazine genes and, thus, pyocyanin production; optimal levels of C_4_HSL are also required. Through a genetic screening approach targeting the RhlR LBP, we recently discovered a RhlR variant (A44M) that could not be activated by C_4_HSL but still maintained its interaction with PqsE and could still bind to DNA [45]; however, gene expression levels of RhlR-regulated genes in a *P. aeruginosa* strain expressing the RhlR A44M variant were similar to that of a 11*rhlR* strain, indicating that, generally, C_4_HSL is required for RhlR transcriptional activation. To further test this in isolates from pwCF, we performed site-directed mutagenesis on the *rhlR* locus to allow for expression of the RhlR A44M variant and then measured pyocyanin levels. Going forward, we use strains 1010 and 1011 due to their ability to produce high levels of pyocyanin, and because we were interested in testing the RhlR dependencies that drive the behaviors to induce pathogenesis (**Figure 3A**). Pyocyanin is one of the primary causes of host cell cytotoxicity during an infection. Clinical isolates expressing the RhlR A44M variant produced significantly less pyocyanin than the parental isolate and to similar levels as the mutant *pqsE^NI^* strain for both sets of isolates (**Figure 3A**). Strains expressing both PqsE and RhlR variants exhibited similar decreases in pyocyanin production, indicating that disrupting just one of the key interactions for RhlR was sufficient to abrogate pyocyanin production (**Figure 3A**). Thus, our *pqsE* and *rhlR* mutant data are consistent with previous studies that identified both the PqsE and the C_4_HSL interaction with RhlR to be necessary for pyocyanin production in laboratory strains. Conversely, we discovered a hypersensitive RhlR variant at position T58 in the LBP in the same genetic screen that identified RhlR A44M as being insensitive to C_4_HSL [45]. The RhlR T58L variant was activated by lower concentrations of C_4_HSL than WT in both *E. coli* and *P. aeruginosa* reporter systems. A *P. aeruginosa* strain expressing RhlR T58L produced less pyocyanin, which we showed was directly due to the hyperactivation of RhlR to repress the *pqsABCDE* operon, thereby reducing PqsE levels to drive RhlR to the phenazine promoters. We observed a similar decrease in clinical isolates that expressed the RhlR T58L variant, indicating that these strains are also sensitive to the concentrations of and subsequent response to C_4_HSL (**Figure 3A**). Lastly, during the initial investigations that characterized PqsE function, it was noted that the enzymatic activity of PqsE is dispensable for its regulation of RhlR-dependent QS traits. To confirm that this was also the case in clinical isolates obtained from pwCF, we introduced the *pqsE* D73A mutation on the chromosome to render PqsE catalytically inactive. Further confirming findings from laboratory strains, the clinical isolates expressing the PqsE D73A variant produced similar levels of pyocyanin as the parental WT strain (**Figure 3A**). In total, these data support the evolutionary conservation of the RhlR:C_4_HSL-PqsE interaction pathways in driving QS signaling in clinical isolates of *P. aeruginosa*.

**Figure 3.**
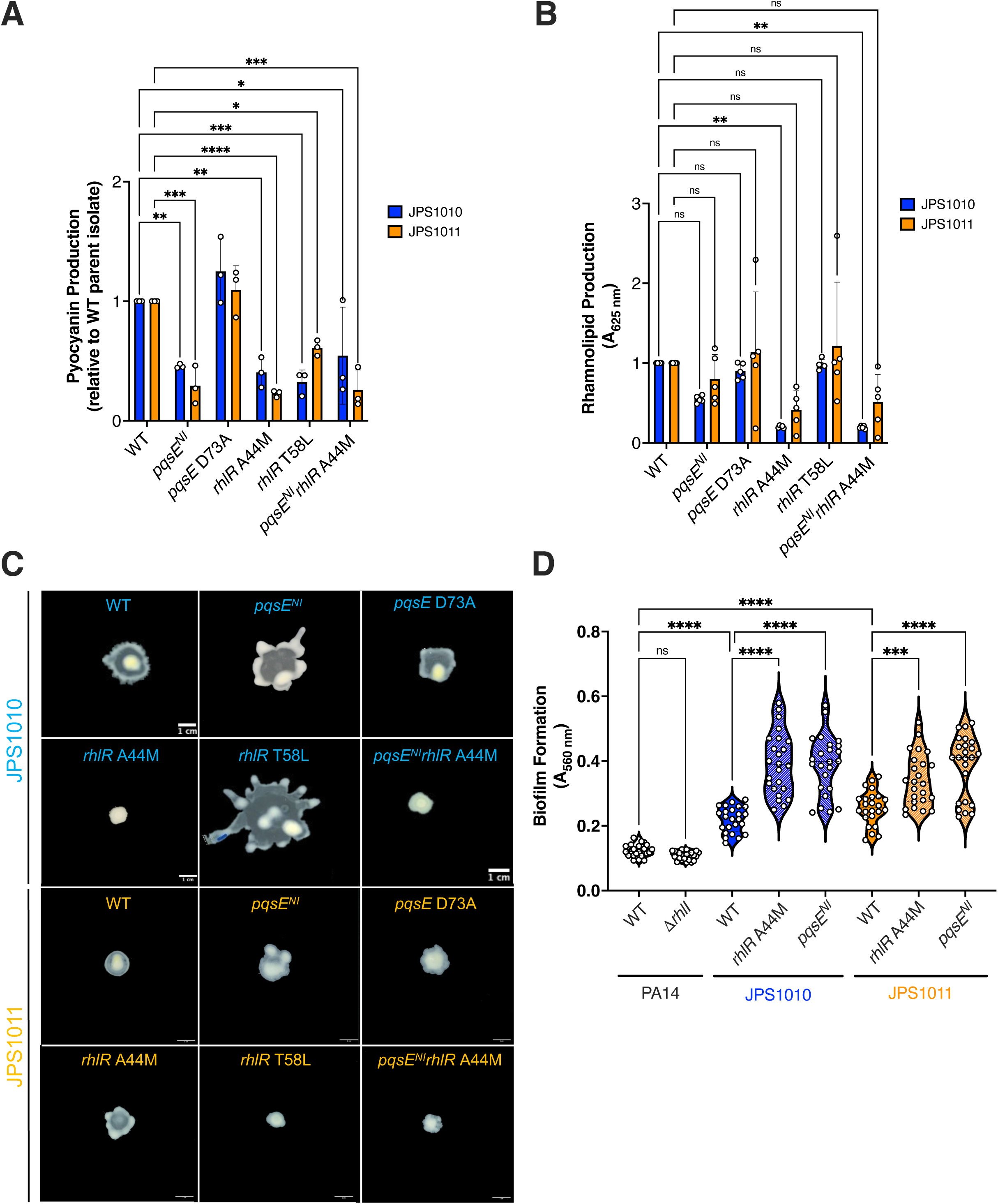
PqsE and C_4_HSL binding contribute to pyocyanin production, rhamnolipid production, and biofilm formation in clinical isolates in a manner like laboratory strains. **A.** Pyocyanin production as measured by OD_695 nm_ / OD_600 nm_ for JPS1010 (blue) and JPS1011 (orange) strains and their corresponding strains containing the labeled *pqsE* and *rhlR* point mutations. All data were normalized to the respective WT parental isolate strain. **B.** Rhamnolipid production as measured by OD _626 nm_ via Victoria Blue PO assays for the same strains as in (**A**). All data were normalized to the respective WT parental isolate strain. **C.** Representative swarming images for WT JPS1010 (blue) and WT JPS1011 (orange) as well their isogenic strains containing the labeled *pqsE* and *rhlR* point mutations that were grown on 0.4% swarming agar. **D.** Biofilm formation of WT PA14 and its isogenic 11*rhlI* mutant strain relative to the JPS1010 (blue) and JPS 1011 (orange) strains and their corresponding strains containing the labeled *pqsE* and *rhlR* point mutations. Statistics were performed using a 2-way analysis of variance (ANOVA) with a Dunnett’s multiple comparisons test. p-value summary: * < 0.05; ** < 0.01; *** < 0.001; **** < 0.0001; ns = not significant.

### RhlR sensitivity to C_4_HSL controls rhamnolipid production in isolates from pwCF

Rhamnolipid production is important for establishing and maintaining infections [81, 82]. Rhamnolipids function as a surfactant for swarming motility across surfaces, including in the host lung [83, 84]. In laboratory strains, the production of rhamnolipids via RhlR-dependent regulation of the *rhlAB* synthases is C_4_HSL-dependent with minimal requirement for PqsE co-regulation [30, 45, 79, 85, 86]. Indeed, clinical isolates followed a similar trend. The disruption of the PqsE dimerization interface led to a small and insignificant decrease in rhamnolipid production, as determined by the absorbance of Victoria Blue PO with cell-free supernatant (**Figure 3B**). The strain that expressed the RhlR A44M variant in the JPS1010 background displayed significantly reduced rhamnolipid production due to the inability of RhlR to respond to C_4_HSL, while the same variant in the JPS1011 displayed a modest but insignificant decline in rhamnolipid production. In contrast to the observed pyocyanin production phenotype (**Figure 3A**), strains expressing RhlR T58L did not exhibit a decrease in rhamnolipid production and, instead, maintained levels comparable to that of the parent strain (**Figure 3B**). Given that the clinical isolates could produce rhamnolipids in a C_4_HSL-dependent manner, we next assessed the ability of the strains to swarm on low percentage agar plates. Interestingly, the parental WT isolates did not exhibit the stereotypical radial swarming that are typically observed in laboratory strains [45] (**Figure 3C**) and only the JPS1010 parental isolate exhibited an expanded area from the initial inoculum, indicative of some motility (**Figure 3C**). Indeed, JPS1011 contained frameshift mutations in *flgE*, *fleQ*, and *motC*, which encode the flagellar hook protein, the major regulator of flagellar gene expression, and a major part of the stator complex that powers the flagellar motor (**Table 1**). Combined, these likely resulting in its inability to use rhamnolipids as a surfactant for colony spread and motility. Additionally, JPS1010 contains many SNPs in motility genes across its genome, although not to the extent of the frameshift mutations observed in JPS1011, and we surmise that this has some effect on motility and the use of rhamnolipids as a surfactant (**Table 1**). Consistent with the rhamnolipid measurements (**Figure 3B**), the JPS1010 mutants containing *pqsE^NI^* and *pqsE* D73A did not exhibit a defect in colony spread (**Figure 3C**), whereas the *rhlR* A44M exhibited a dramatic decrease in colony spread (**Figure 3C**). Furthermore, the isolates containing *rhlR* T58L, which maintained increased colony spread (**Figure 3C**). Isolates expressing the double mutation of *pqsE^NI^*and *rhlR* A44M phenocopied a *rhlR* A44M, indicating that C_4_HSL was the dominant driver of rhamnolipid production and colony spreading. The different strains with their isogenic mutants expressed RhlR to similar levels across backgrounds (**Figure S2**). In total, these data support that, like laboratory strains, RhlR sensitivity to C_4_HSL is important for rhamnolipid production and swarming in clinical isolates.

### PqsE and C_4_HSL repress biofilm formation in clinical isolates from pwCF

Biofilm formation is a hallmark of chronic infections and a primary driver of antibiotic tolerance that complicates treating *P. aeruginosa* infections [17, 22, 24, 87, 88]. To determine the role of QS in regulating biofilm formation in clinical isolates, we performed static biofilm growth experiments and used crystal violet staining as a measurement for biofilm attachment to surfaces. To the best of our knowledge, there are no published studies that investigate the role of the Rhl system on pellicle formation in static biofilms. However, much more is known about the role of the Rhl-PqsE module in colony biofilm formation, which established the current paradigm that RhlR and PqsE function as repressors of biofilm traits; the deletion of *rhlR* and *pqsE* led to hyper-rugose colony biofilms. Meanwhile, C_4_HSL is involved in biofilm in an opposing mechanism, as deletion of *rhlI* led to smooth colony biofilms, via a mechanism that is not yet understood. In the laboratory strain, the RhlI system had no effect on static biofilm growth, as both WT and 11*rhlI* strains had similar crystal violet measurements (**Figure 3D**). As expected from chronic infection isolates, the parental strains of JPS1010 and JPS1011 had significantly higher levels of adhesion to surfaces and enhanced pellicle formation compared to the laboratory strain. The biofilm phenotypes in both isolates were enhanced by disrupting QS, either in disrupting the activation of RhlR by C_4_HSL or PqsE (**Figure 3D**), indicating that RhlR functions as a repressor of biofilm traits in clinical isolates during static growth conditions, which is consistent with its previously characterized role in colony biofilms formed by laboratory strains [57, 76–78].

### PqsE and C_4_HSL co-regulate RhlR-dependent QS gene expression in isolates from pwCF

To determine the totality of the regulatory impact PqsE and C_4_HSL has on RhlR signaling and to establish a core regulon for these inputs in clinical isolates, we performed RNA-seq experiments to compare the JPS1010 and JPS1011 parent CF isolates with their respective *rhlR* A44M and *pqsE^NI^* mutants (**Figures 4A-D**, **Table 2**, **Table S4**). Initially, we compared the differential expression of genes within an isolate to determine the role of the C_4_HSL- (**Figures 4A** and **4B**, **Table 2**) and PqsE-dependent regulon (**Figure 4C** and **4D**, **Table 2**). In total, JPS1010 and JPS1011 differentially expressed 470 and 591 genes in a C_4_HSL- and/or PqsE-dependent manner, respectively (**Figure 4E**, **Table 2).** JPS1010 and JPS1011 had 33 and 50 of the total differentially regulated genes that were shared between both RhlR activators, respectively (**Figure 4E**, **Table 2**).

**Figure 4.**
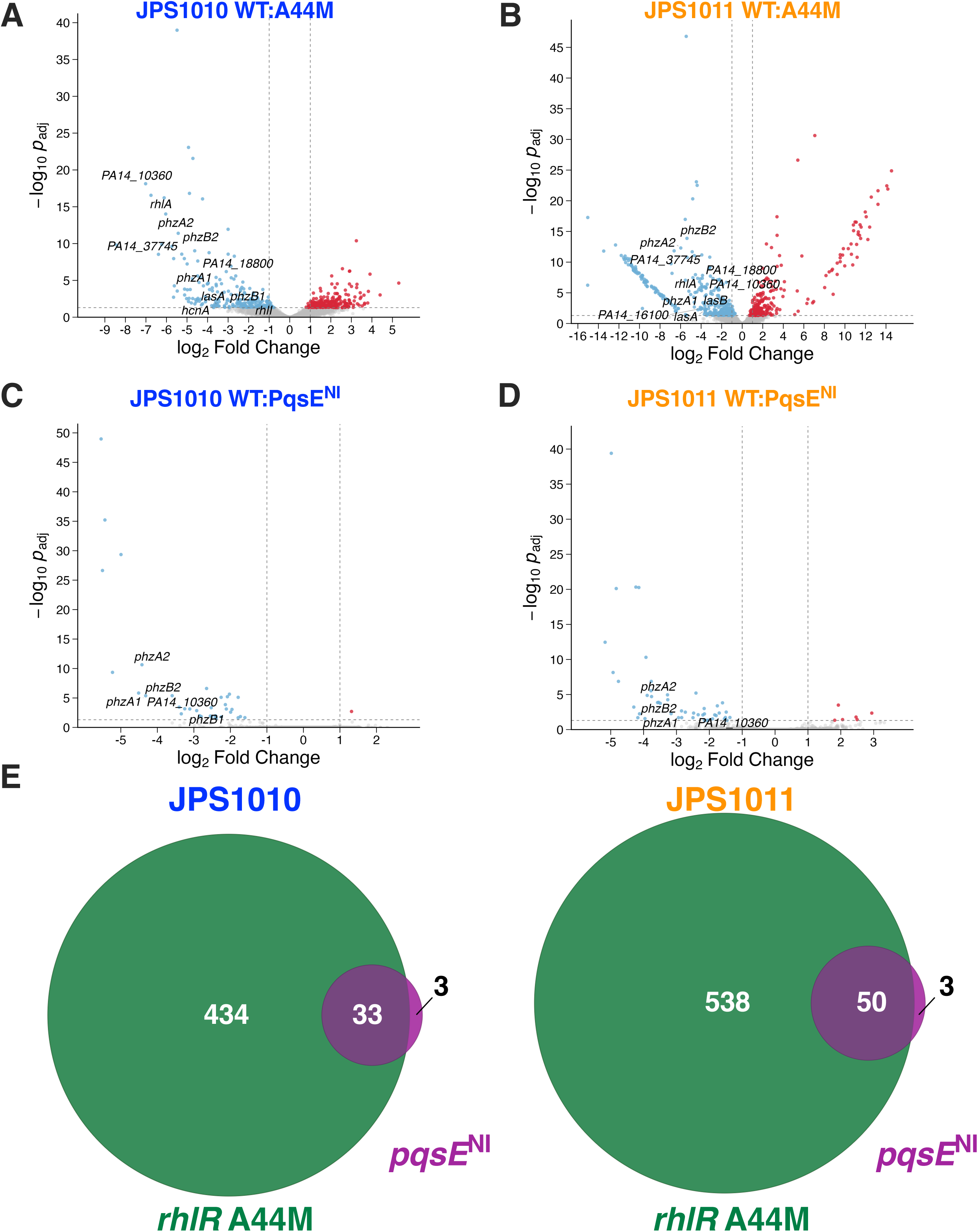
PqsE and C_4_HSL binding have different contributions to RhlR-dependent gene expression. Volcano plots depicting the differential gene expression of **A.** WT JPS1010 and **B.** WT JPS1011 compared to their corresponding *rhlR* A44M mutant strains. Volcano plots depicting the differential gene expression of **C.** WT JPS1010 and **D.** WT JPS1011 compared to their corresponding *pqsE^NI^* mutant strains. All genes that are labeled were considered differentially expressed and known to be directly regulated by RhlR via previous ChIP-seq analyses [79]. **E.** Venn diagrams depicting the overlap in differentially regulated genes for JPS1010 *rhlR* A44M and *pqsE^NI^* (left) and JPS1011 *rhlR* A44M and *pqsE^NI^* (right).

**Table 2.** Differentially expressed genes in JPS1010 and JPS1011 mutant strains.

The C_4_HSL-dependent regulon was largely conserved between JPS1010 and JPS1011. There was an overlap of 199 genes that were differentially regulated in a C_4_HSL-dependent manner in JPS1010 and JPS1011, representing 43% and 34% of the total C_4_HSL-dependent genes in the respective isolates (**Figure 5A**, **Table 2**). Several of these genes are well-characterized from laboratory strains as being C_4_HSL-dependent, namely genes involved in hydrogen cyanide production (*hcn*), phenazine production (*phz*), rhamnolipid production (*rhl*), quinolone production (*pqs*), protease production (*las*), and chitin degradation (*chiC*) (**Table 2**). Interestingly, consistent with the rhamnolipid production phenotype (**Figure 3B**), the *rhlR* A44M mutant had a more dramatic effect on *rhlA* gene expression in JPS1010 (log_2_FC = -6.8) than JPS1011 (log_2_FC = -4.1) (**Table 2**). Meanwhile, genes in the phenazine operon displayed more consistent downregulation across the two isolates: *phzC1* was downregulated by a log_2_FC of -4.9 and -4.0 in JPS1010 and JPS1011, respectively (**Table 2**). These nuances in shared RhlR:C_4_HSL-regulated genes highlight the conserved core of RhlR-dependent signaling in distantly-related clinical isolates. Conversely, gene expression levels that were elevated in the backgrounds expressing A44M presented unique insights into the repressive function of RhlR-C_4_HSL. Indeed, many of these genes have not been previously characterized as being dependent on QS. For example, the *phoPQ* and *oprH* genes (log_2_FC = +2.4, +1.9, and +2.9, respectively) involved in magnesium signal transduction, the *trpAB* genes (log_2_FC = +2.4 and +3.2, respectively) involved in tryptophan biosynthesis, the *bfiSR* genes (log_2_FC = +1.6 and +2.0, respectively) involved in biofilm formation, and the *prtN* (log_2_FC = +2.9) gene responsible for activating R-type pyocins were all significantly elevated when RhlR-C_4_HSL signaling was disrupted (**Table 2**). Additionally, there are hundreds of uniquely C_4_HSL-dependent genes in the two different strains, many of which are uncharacterized (**Table 2**). The existence of isolate-specific RhlR-dependent gene regulation likely represents the ability of certain isolates to evolve different QS-dependent regulons, a reflection of the adaptation of the isolate to its environment.

**Figure 5.**
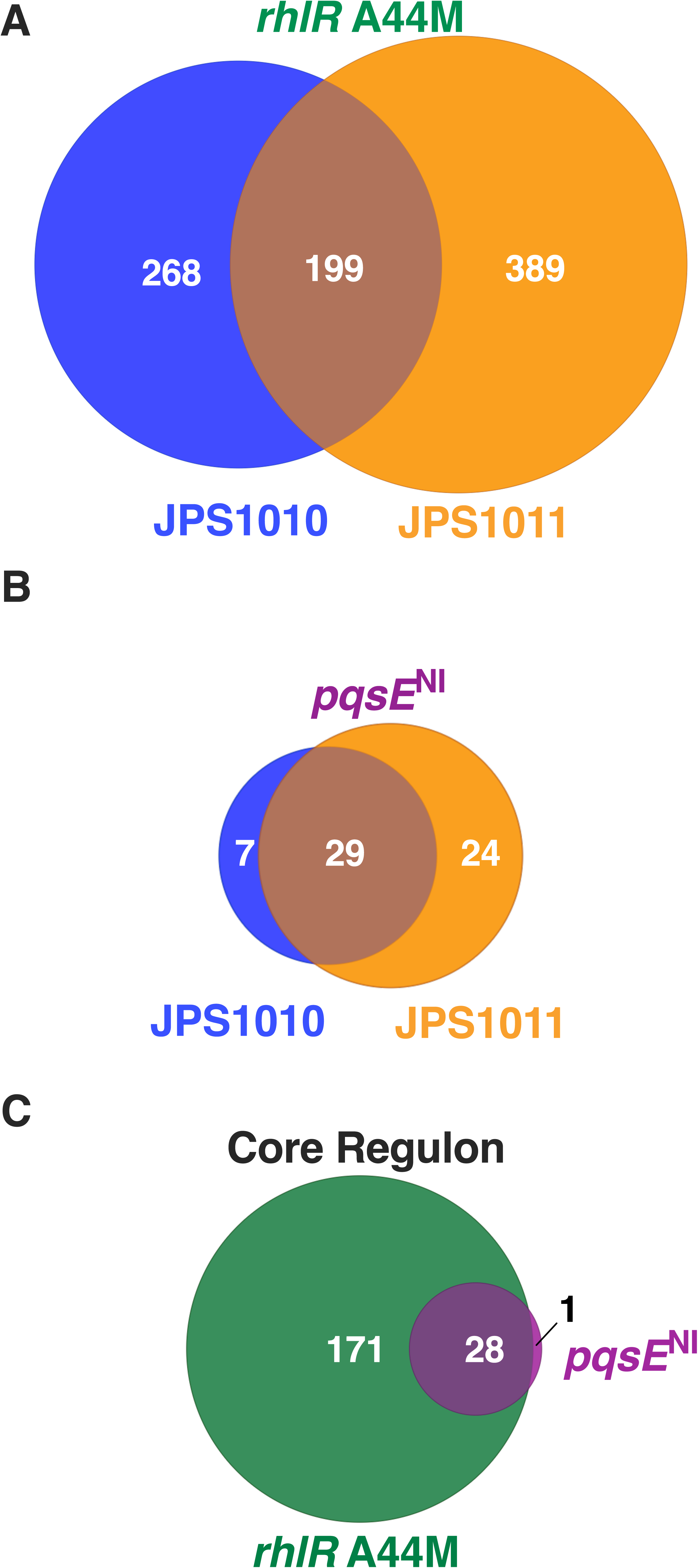
The RhlR core regulon in clinical isolates consists of 28 genes. **A.** Venn diagram depicting the overlap in differentially regulated C_4_HSL-dependent genes for JPS1010 (blue) and JPS1011 (orange). **B.** Same as in (**A**) except for PqsE-dependent genes. **C.** Venn diagram depicting the RhlR core regulon as gleaned from the overlap shown in (**A**) and (**B**).

Similar to the overlap in C_4_HSL-dependency, there was an overlap of 29 genes that were differentially regulated in a PqsE-dependent manner in JPS1010 and JPS1011, representing 81% and 55% of the total PqsE-dependent genes in the respective isolates (**Figure 5B**, **Table 2**). The genes shared between *pqsE^NI^* mutants in JPS1010 and JPS1011 backgrounds include genes that encode canonical virulence factors such as the phenazines (*phz*), phenazine modifying enzymes (*pumA*), the efflux system responsible for expelling phenazines (*mexGHI*), iron-sulfur-binding clusters (*iscAU* and *hscAB*), and nitrate detoxifying enzymes that support biofilm growth and/or growth in anaerobic environments (*ahpF* and *trxB*). Thus, like C_4_HSL, PqsE exerts well-conserved control over phenazine production and the associated oxidative stress response pathways to manage the redox consequences of that production in clinical isolates. Conversely, the clinical isolates revealed disparate RhlR-PqsE-dependent control over *hcnA* and *lecB* expression, two genes that were previously implicated as being dependent on PqsE for RhlR promoter binding and gene expression [79]. Specifically, the expression of *hcnA* and *lecB* decreased significantly in the *pqsE^NI^*mutant background of JPS1011 with a log_2_fold reduction of -3.2 and -2.9, respectively. Meanwhile, the *pqsE^NI^* mutant background of JPS1010 exhibited a log_2_fold reduction in expression of -0.3 and -1.0 for *hcnA* and *lecB*, respectively, which were deemed insignificant by our analyses. These data demonstrate that while PqsE-dependent control of phenazine biosynthesis is conserved, the PqsE-RhlR module can differentially regulate other pathways, including hydrogen cyanide and lectin production. Thus, isolate specific variation exists even within the well-conserved RhlR-C_4_HSL-PqsE QS module, highlighting the complexity of the biology underlying RhlR signaling, especially within the context of CF-adapted strains.

Of the 199 C_4_HSL-dependent and 29 PqsE-dependent differentially expressed genes, 28 of them were shared between the isolates and their genotypes, representing a core RhlR-dependent regulon (**Figure 5C**, **Table 2**). This core regulon defines the most robustly regulated outputs of the RhlR transcriptional network in the context of clinical isolates from pwCF and includes the phenazines and MexGHI-OpmD efflux system among other pathways related to oxidative stress defense and iron-sulfur cluster biogenesis, encompassing both well-characterized and uncharacterized genes. Thus, we hypothesize that conservation across both RhlR inputs and isolate backgrounds indicates that these genes are central to RhlR function in the context of infection.

### RhlR-dependent pathogenesis is more dependent on C_4_HSL than PqsE

To determine the role of the individual dependencies for RhlR-dependent signaling in an infection model, we used a mammalian cell culture infection model. Confluent Calu-3 lung epithelial cells were infected with different *P. aeruginosa* isolates, their isogenic QS mutant strain, or the lab strain PA14 at an MOI of 10. Subsequently, the media was harvested for cytokine analysis via cytometric bead array analysis using fluorescent sorting. We hypothesized that clinical isolates obtained from pwCF would elicit a pro-inflammatory cytokine response in airway epithelial cells due to isolate-specific changes of bacterial surface markers and virulence factor gene expression. Consistent with their status as pathogenic clinical isolates, JPS1010 and JPS1011 induced higher levels of IL-8 cytokine production than WT PA14 (**Figure 6A**). These elevated levels of IL-8 production were significantly reduced in all QS mutant backgrounds for both the JPS1010 and JPS1011 strains. Across the panel of IL-8 (**Figure 6A**), IL-6 (**Figure 6B**), and TNF (**Figure 6C**), JPS1011 induced elevated levels of cytokine production, with the TNF production profile being unique to JPS1011 pathogenesis, as neither PA14 nor JPS1010 induced TNF production. Expectedly, deletion of *rhlR* had the most pronounced effect on all cytokine production, regardless of strain background (**Figure 6A-C**). Meanwhile, strains expressing *pqsE^NI^* exhibited intermediate phenotypes for IL-8 and IL-6. Conversely, strains expressing *pqsE^NI^* had no effect on JPS1011 induction of TNF production. Strains expressing *rhlR* A44M induced cytokine production levels like a 11*rhlR* strain, especially in the case of both IL-6 and TNF. These data are consistent with our transcriptional profiling and phenotypic assays, indicating that the RhlR-C_4_HSL interaction is a key driver of the proinflammatory response in epithelial cells. Interestingly, a strain expressing both *pqsE^NI^* and *rhlR* A44M did not match the results of either the 11*rhlR* or *rhlR* A44M individual mutants for any of the cytokine production profiles (**Figure 6A-C**). Instead, these strains induced cytokine production profiles like strains expressing *pqsE^NI^*, indicating that RhlR-dependent signaling in the absence of C_4_HSL and the PqsE interaction can drive some pathogenic traits, perhaps through the elevated levels of genes involved in acute infection pathways.

**Figure 6.**
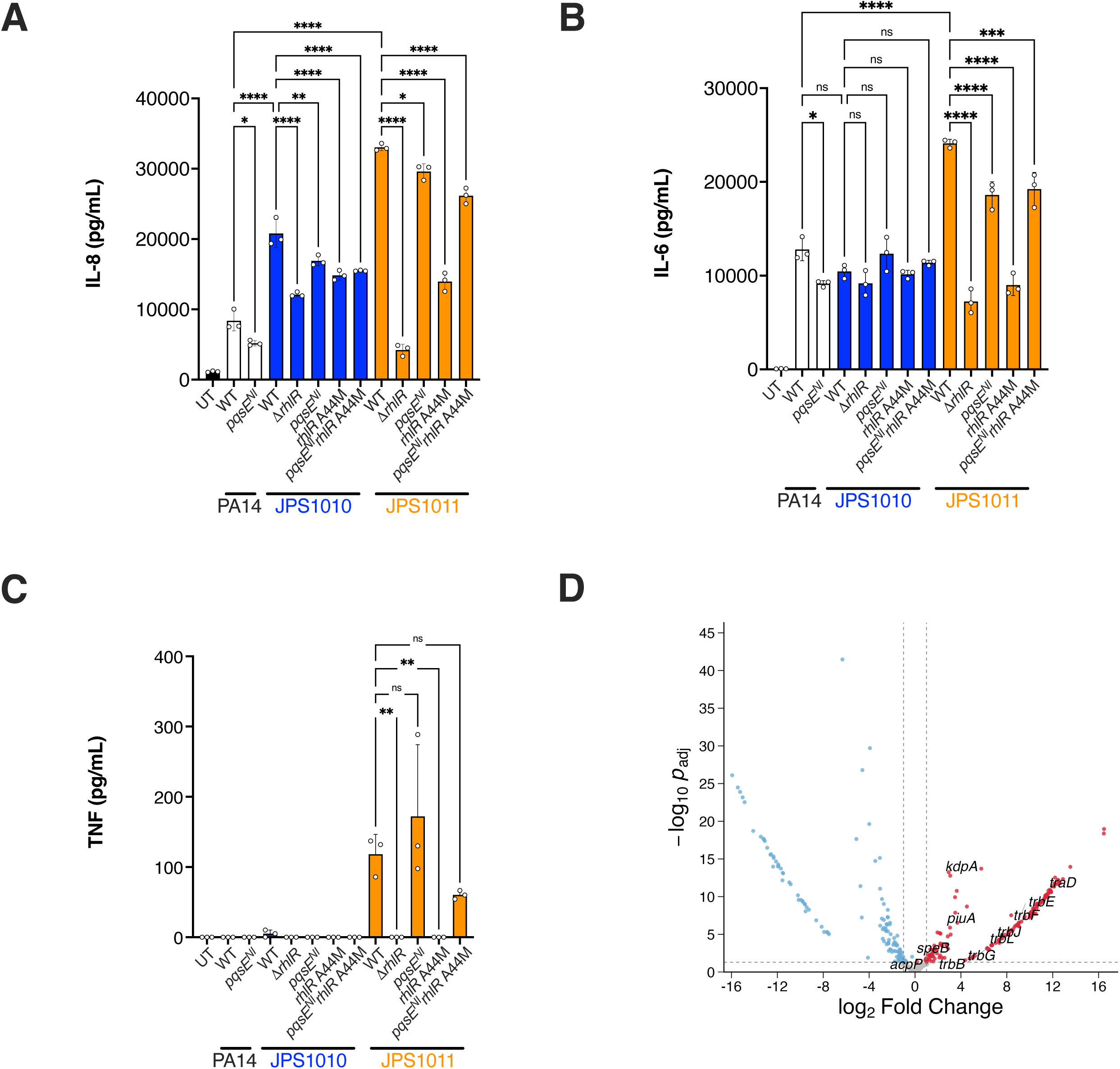
RhlR and its conserved PqsE- and C_4_HSL-dependent regulon are required for pathogenesis in a mammalian cell culture infection model. **A.** IL-8, **B.** IL-6, and **C.** TNF cytokine production by Calu-3 cells in response to the labeled WT PA14 (white), JPS1010 (blue), and JPS1011 (orange) strains as measured by human inflammatory cytokine cytometric bead array (CBA) I kit. Statistics were performed using a 2-way analysis of variance (ANOVA) with a Tukey’s multiple comparisons test. p-value summary: * < 0.05; ** < 0.01; *** < 0.001; **** < 0.0001; ns = not significant. **D.** Volcano plots depicting the differential gene expression of WT JPS1010 and WT JPS1011. Positive values indicate genes that are upregulated in JPS1011 relative to JPS1010; negative values indicate genes that are downregulated in JPS1011 relative to JPS1010 (*i.e.*, upregulated in JPS1010 relative to JPS1011).

To determine the molecular basis for the increased TNF production by epithelial cells in the presence of JPS1011 but not JPS1010, we compared the transcriptional profiles of the WT parental strains of both clinical isolates (**Figure 6D**, **Table 3**). Each strain exhibited distinct transcriptional profiles outside of their core QS functions. Given the genomic diversity between JPS1010 and JPS1011, it was difficult to ascertain which genes are specifically differentially regulated at the level of transcription because of adaptation to the host lung to cause TNF induction or are merely the result of divergent evolution. A few candidate upregulated genes that appear to be the result of adaptation to the host stood out as playing a role in TNF production by epithelial cells and the subsequent response by *P. aeruginosa*: *acpP, piuA*, *trb*/*traD*, and *kdpA*. *acpP* encodes for an acyl carrier protein that donates fatty acid chains to lipid A, which is a key driver of TNF induction in human cell lines [89–92]; *piuA* encodes for a TonB-dependent iron uptake receptor that is linked to persistence in the airway of pwCF [93, 94]; *trbBEFGJL* and *traJ* encode for type IV secretion/conjugation apparatus, which can induce MAP kinases to increase the proinflammatory cytokine response [95, 96]; *kdpA* encodes for a potassium transporter that is involved in the uptake of potassium secreted by airway epithelial cells [97]. Collectively, expression of these genes indicates that JPS1011 has specifically evolved a signaling system tuned to the host lung that can drive strain-specific induction of TNF.

**Table 3.** Differentially expressed genes between JPS1010 and JPS1011 parental strains.

## DISCUSSION

The role of RhlR signaling in clinical isolates from pwCF had been limited to full gene deletions of *rhlR* [75, 77]. These studies established the foundation that RhlR is the central QS transcriptional regulator in CF isolates. Given the nuanced role of C_4_HSL and PqsE in activating RhlR, we directly assessed the specific contribution of each regulator to RhlR function by using variants of RhlR that disrupt its ability to be activated by C_4_HSL as well as variants of PqsE that disrupt its ability to interact with RhlR [45, 62]. Primarily, our separation-of-function point mutations revealed that prior analyses of a core regulon is a composite of both C_4_HSL- and PqsE-dependent sub-regulons. For example, prior work showed that full deletion of *rhlR* revealed that *hcnA* and *lecB* were members of the core regulon; here, we show that *hcnA* and *lecB* show isolate-specific PqsE-dependency. Given the complexities of the interplay between C_4_HSL and PqsE levels, we suspect that certain important biological processes that are differentially regulated by the two different inputs would obscure interesting regulatory connections. Here, we showed that the PqsE-RhlR interaction plays a critical role to regulate oxidative stress responses (*i.e.*, *ahpF*, *trxB*, and *katB*, which are enzymes involved in the peroxide detoxication pathway, thiol redox homeostasis pathway, and catalase pathway, respectively) and iron-sulfur cluster assembly (*i.e., iscA*, *iscU*, *hscA,* and *hscB)* in clinical isolates from pwCF, while the RhlR-C_4_HSL interaction plays a similarly important role to regulate two-component signal transduction systems (PhoPQ), both of which have not been previously characterized as core genes or pathways tied to the RhlR-C_4_HSL-PqsE module.

A recent analysis of PqsE in twelve clinical isolates revealed that while its sequence is highly conserved, the deletion of *pqsE* can have variable effects on several different phenotypic outputs; a *pqsE* deletion resulted in strain-specific patterns of phenazine and rhamnolipid production compared to their respective parent strains [80]. Thus, there are aspects to RhlR-PqsE signaling that we do not yet fully understand. However, our work here confirms these findings with the added layer of specifically dissecting the RhlR-PqsE interaction by introducing the *pqsE^NI^*mutation on the chromosome and by performing transcriptomics, which revealed an expanded set of differentially expressed genes, including genes that were not previously characterized to be part of the RhlR-PqsE regulon.

Our separation-of-function mutations, specifically the use of *rhlR* A44M, revealed an expanded regulon containing genes normally repressed by RhlR-C_4_HSL that is absent from the strains expressing *pqsE* mutants. Our transcriptomic analyses revealed that RhlR-C_4_HSL functions as an indirect repressor of *phoPQ*, *bfiR*, and *prtN* in clinical isolates obtained from pwCF. It is intriguing to speculate why RhlR might normally function as a repressor of these genes in a clinical isolate from a chronic infection. PhoPQ signal transduction, BfiR-mediated biofilm initiation, and PrtN-mediated interspecies competition are all pathways that might be required during early colonization, early infection, or acute infection but not during chronic infection [12, 98–103]. Thus, RhlR-mediated repression of these traits in parallel with RhlR-dependent positive regulation of phenazine production and detoxification is consistent with RhlR as a central regulator of virulence traits and pathogenesis during both acute and chronic infection.

The observation that the *pqsE^NI^ rhlR* A44M double mutant induces cytokine production profiles like the *pqsE^NI^*mutants rather than the complete disruption observed by strains expressing *rhlR* A44M alone was an unexpected finding but suggests nuances about RhlR-dependent signaling that have not been previously appreciated. We believe this result can be interpreted in different ways. First, it suggests that RhlR-dependent signaling in the absence of both co-regulatory inputs can proceed with RhlR alone. This is without precedent in the current literature on RhlR function, but it cannot be ruled out based on our data. Second, the de-repression of multiple pathways in the absence of both regulators maximally drives acute infection that now initiate a similar cytokine response in epithelial cells as the WT parental isolate. Third, PqsE^NI^ could exert RhlR-independent effects on bacterial physiology in clinical isolates that can only be observed when RhlR-C_4_HSL function is disrupted. Again, this is currently without precedence in the literature and requires future investigation. Regardless of the mechanism, this finding highlights the complexity of RhlR co-regulator interactions in the context of host infection and underscores that the consequences of simultaneous disruption of the two inputs on RhlR signaling may be difficult to predict. Indeed, based on our infection model data, it appears that disrupting RhlR function through its LBP might be sufficient to reduce *P. aeruginosa* pathogenesis. Taken together, these findings underscore the need to evaluate anti-QS therapeutic strategies and, indeed, basic mechanisms of pathogenesis, in clinically relevant isolate backgrounds and across multiple infection model systems before advancing such approaches toward therapeutic development.

Initially, it was surprising to us that JPS1011 induced such a strong TNF cytokine response. Bacteria can modulate TNF function indirectly via alteration of upstream signaling pathways in the host such as NF-κB and MAPK, likely through inactivating proteins or changes to the bacterial cell surface (*i.e.*, changes to LPS or the loss of flagella/pili; canonical bacterial PAMPs) [104]. Indeed, many Gram-negative bacteria such as *Yersinia* species, *Salmonella typhimurium*, *Escherichia coli* K1 and STEC, *Bordotella bronchoseptica*, *Neisseria gonorrhoeae*, *Bartonella henselae*, and *Brucella suis* all produce proteins that result in the downregulation of TNF production [104]. It was shown previously that the loss of flagellin [105] or increased HHQ/PQS signaling [106] can downregulate NF-κB signaling. The type III secretion system (T3SS) has also been shown to play a role in TNF production; machinery components and the ExoU toxin can activate NF-κB, while the adenylate cyclase ExoY attenuates both NF-κB and MAPK signaling [107]. Our transcriptional profiling of JPS1011 revealed multiple pathways that likely converge on a similar output to these previously proposed mechanisms. Collectively, cytokine profiling experiments indicate that JPS1011 is highly adapted to the host environment; surface engagement, iron acquisition, ligand presentation that triggers the innate immune system, and contact-dependent host cell activation drive a robust TNF response that is absent from other strains, including JPS1010. In total, these data highlight the multiple ways in which *P. aeruginosa* can control QS traits and cause infections.

## MATERIALS AND METHODS

### Molecular biology and strain construction

All plasmids used in this study were previously constructed [45, 62]. All strains and plasmids used in this study are in **Table S5**. Standard cloning and molecular biology techniques were used to generate *P. aeruginosa* clinical isolate mutations. Introduction of genes encoding RhlR and PqsE variants onto the isolate chromosome was achieved using previously published protocols involving the pEXG2 vector. The pEXG2 vector containing RhlR and PqsE variants as well as deletion constructs were transformed into *E. coli* SM10 λ*pir* followed by conjugation into the appropriate clinical isolate. All strains were confirmed by amplifying the target locus followed by Sanger sequencing.

### Whole genome sequencing and bioinformatic analyses

<u>DNA Extraction:</u> Overnight cultures were grown in 3 mL of LB media. Cells were harvested by centrifugation at 16,000 x *g* for 5 min at 4 °C. Supernatant was discarded, and cells were resuspended in 20 μL 1X PBS. 20 μL of Proteinase K and 20 μL of CLE3 buffer (PacBio) was added to each sample and pulse mixed for 10 s with a vortex. Samples were incubated at 55 °C and periodically mixed by inversion for 10 min. 20 μL of RNase A was added and pulse vortexed for 5 s and then incubated at RT for 3 min. 100 μL of Buffer BL3 (PacBio) was added and pulse vortexed for 10 s. Samples were incubated at 70 °C on a heat block and periodically mixed by inversion for 20 min, then pulse vortexed for 10 s. A Nanobind (PacBio) disk was added, then 100 μL of isopropanol and mixed by inversion five times. Samples were mixed on a tube rotator at 9 rpm for 10 min. A magnetic rack was used to stabilize disks and then the supernatant was removed, avoiding the disk. 700 μL of Buffer CW1 was added and mixed by inversion four times. The supernatant was discarded in the same manner. 500μL of Buffer CW2 was added and mixed by inversion four times. The supernatant was discarded in the same manner. Samples were spun for 2 s on a mini-centrifuge, and the residual supernatant was discarded. 200 μL Buffer LTE (PacBio) was added and incubated for 10 min at RT. The eluate was collected in a new tube. Nanobind disks were spun for 15 s at 10,000 x *g* to remove residual eluate. All liquid was combined with for the total eluate. The samples were pipette mixed 10 times and left at RT overnight to fully solubilize. The samples were mixed again, and the concentration was taken using a NanoDrop (ThermoFisher) to ensure a homogeneous sample. Samples were stored at -20 °C until sequencing. <u>Data Analysis:</u> The DNA samples were sequenced using MinIon (Oxford Nanopore). Fastq files generated from sequencing were trimmed using Porechop to identify and remove adapters from the middle and ends of reads [108]. De novo assembly was performed on trimmed sequences using Flye [109]. Variant calling for single nucleotide polymorphisms and insertions or deletions was performed on the assemblies using Snippy [110]. All SNPs were plotted using Circos [111].

### C_4_HSL quantification

AHL concentrations were quantified from cell-free supernatants using a UHPLC-HRMS method as previously described [73]. Briefly, filtered supernatants were extracted in methanol, clarified by centrifugation, and diluted 1:1 in 0.1% formic acid prior to analysis on a Vanquish UHPLC coupled to a QE-Orbitrap mass spectrometer operating in positive ESI mode. Analytes were separated on a C18 column using a methanol/water gradient and quantified against an external calibration curve using a 5 ppm accurate mass window.

### Pyocyanin production

Pyocyanin production assays were performed as previously described [56]. Overnight cultures were diluted 1:100 in 25 mL LB and grown for 6-8 h or until OD_600 nm_ of ∼2.0. 1 mL aliquots were then pelleted at 24,104 x *g* and the supernatant was collected. The absorbance at was read at 695 nm. Pyocyanin production was determined by plotting OD_600 nm_/OD_695 nm_.

### Rhamnolipid production

Clear 96-well plates (CellTreat) were stained with 50 μL of 0.1 mg/ml Victoria Pure Blue BO dye in isopropanol. The isopropanol was evaporated under vacuum for 1 h at 45 °C. 300 μL of 0.5M NaOH was added to the wells and incubated at RT for 10 min and then dried under vacuum for 1 h at 45 °C. 5 mL cultures were grown overnight in LB. 1 mL of overnight cultures were centrifuged for 2 min at 24,104 x *g*. Cell-free culture supernatants were added in triplicate to stained wells and incubated with agitation at 750 rpm for 1 h at RT. 200 μL were transferred to a clean 96-well plate (CellTreat) and the absorbance was read at 625 nm (Molecular Devices SpectraMax M5 SoftMax Pro5).

### Swarming assays

Swarming plates were made with 5X M9 Salts (20 mM NH_4_Cl, 12 mM Na_2_HPO_4_, 22 mM KH_2_PO_4_, 8.6 mM NaCl), 0.4% agar, 0.5% casamino acids, 1 mM CaCl_2_, 1 mM MgSO_4_, and 1.35% (w/v) dextrose. 5 mL of overnight cultures were grown in LB and normalized to an absorbance of 600 nm = 1.0. 5 μl of normalized culture was spotted onto fresh swarming media and left to dry. Plates were incubated face down at 37 °C for 24 h and then at RT for another 12 h. Images of plates were taken using Epson Perfection V850 Pro scanner at a resolution of 300 dpi. <u>Image analysis</u>: Swarming plates were analyzed using our previously published protocol [45].

### Static biofilm

Static biofilm growth and biofilm quantification were adapted from a previous study [112]. Overnight cultures were diluted 1:100 into fresh LB and 100 μL was added to a 96-well plate in replicates of eight. The plate was incubated statically at 37°C for 24 hours. Cells were removed from the plate, and the plate was gently washed to remove excess cells and media. 125 μL of 0.1% crystal violet in water was added to each well and incubated for 15 minutes. Crystal violet was removed, and the plate was washed thoroughly and allowed to dry for several hours or overnight. 125 μL of 30% acetic acid was added to each well and incubated for 15 minutes. The solubilized crystal violet was transferred to a fresh 96-well plate and absorbance was read at 560nm with a 30% acetic acid blank.

### RNA extraction, sequencing, and differential gene expression analyses

<u>RNA extraction</u>: 5mL Overnight cultures were grown shaking at 37 °C. Cells were harvested by centrifugation at 500 x *g* for 5 min at 4 °C. Culture supernatant was discarded, and the pellets were frozen until RNA extraction. RNA was extracted using Invitrogen PureLink RNA Extraction Mini Kit (Invitrogen). Cell pellets were resuspended in 100 μL of lysozyme solution containing 10 mM Tris-HCl pH 8.0, 0.1 mM EDTA, and 1 mg/mL lysozyme. 5 μl of 1% SDS was added, and the solution was mixed with a vortex, and then incubated at RT for 5 min. Resuspended cells were lysed by bead beating (BioSpec 0.1 MM Zirconia/Silica Beads) two times for 50 s each. Beads were allowed to settle, then the lysed samples were transferred to a clean tube. 250 μL of 100% ethanol was added to each sample and mixed with a vortex. The samples were transferred to a spin cartridge (Invitrogen), and passed through the filer at 12,000 x *g* for 15 s. The flowthrough was discarded. 700 μL of Wash Buffer I (Invitrogen) was added, and samples were centrifuged at 12,000 x *g* for 15 s. The flowthrough was discarded. 500 μL of Wash Buffer II (Invitrogen) was added, and samples were centrifuged at 12,000 x *g* for 15 s. Flowthrough was discarded. The spin cartridges were dried by centrifugation at 12,000 x *g* for 1 min. 50 μL RNase-free water was added to the cartridges and incubated at RT for 1 min. Samples were then eluted into a new collection tube by centrifugation at >12,000 x *g* for 2 min. The RNA extractions were flash frozen using liquid nitrogen and stored at -80 °C until sequencing. <u>Data analysis:</u> Bulk RNA was sent to SeqCenter (Pittsburgh, PA) where it was sequenced with paired-end Illumina sequencing with rRNA depletion using Ribo-Zero Plus (Illumina). Quality of reads was checked using FastQC [113] and reads were mapped to the *Pseudomonas aeruginosa* UCBPP-PA14 reference genome using Bowtie2 [114]. FeatureCounts was used to count mapped reads [115]. Differential expression was determined using DESeq2 to make pairwise comparisons using the Wald test [116]. Volcano plots were created using ggplot2 package in R [117].

### Mammalian cell infection assay

Calu-3 cells (ATCC, #HTB-55) were maintained in EMEM supplemented 10% FBS, 1% L-glutamine, and 1% penicillin/streptomycin at 37 °C with 5% CO_2_. For cytokine detection, cells were seeded in 6-well dishes at 5×10^5^ cells per well and allowed to grow to confluency. Overnight cultures of *P. aeruginosa* were pelleted and washed three times in DPBS. Cells were normalized to an OD_600_ of 0.7 in cell culture media. *P. aeruginosa* was allowed to attach to the epithelium for 2 h and Calu-3 cells were subsequently washed three times in DPBS to remove any unattached bacteria. Infection was allowed to proceed for an additional 4 h, at which time spent media was collected and stored at -80 °C. CFU were enumerated to ensure similar growth across *P. aeruginosa* strains tested. The human inflammatory cytokine cytometric bead array (CBA) I kit (BD Biosciences) was used to determine the amount of IL-12p70, TNF, IL-10, IL-6, IL-1ß, and IL-8 present. Samples were prepared according to manufacturer’s instructions. Samples were read using a FACSymphony A3 Cell Analyzer (BD Biosciences).

### Western blot analyses

Cells were harvested by centrifugation at 21,300 x *g* for 2 min and pellets were immediately resuspended in 2X SDS sample buffer normalized to OD_600 nm_ and then flash frozen in liquid nitrogen for storage at -20 °C. Samples were boiled at 100 °C for 5-10 min, pelleted at 21,300 x *g* for 2 min, and 10 µL of supernatant was resolved on precast SDS-PAGE gels (Bio-Rad) at 35 mA for 35 min. Protein was transferred to PVDF membrane (Bio-Rad) at 110 A for 1 h using a semi-dry transfer cell (Bio-Rad) and blocked with 5% milk in TBST for 2 h at RT. Membranes were incubated overnight at 4 °C with a polyclonal rabbit α-RhlR antibody (Cambridge Antibodies) at 1:1,000 dilution, washed 3X with TBST (5 min each), and incubated with HRP-conjugated goat anti-rabbit secondary antibody (Thermo Fisher Scientific) at 1:10,000 for 1 h at RT. After three 15 min washes with TBST, blots were developed with Pierce ECL substrate (Thermo Fisher Scientific) and imaged with a 2-minute exposure on an iBright-1500 (Thermo Fisher Scientific).

## DATA AVAILABILITY

Sequencing data have been deposited at the NCBI Sequence Read Archive under the submission number SUB16149246 (Bioproject # PRJNA1460534).

## SUPPORTING INFORMATION

**Table S1. Comparative genomics of clinical isolates relative to PA14.**

**Table S2. PA14 genomic regions absent or divergent in CF Clinical Isolates.**

**Table S3. Novel genomic regions in clinical isolates relative to PA14.**

**Table S4. Genome-wide transcript levels from RNA-seq.**

**Table S5. Strains and plasmids used in this study.**

---

**Figure S1.**
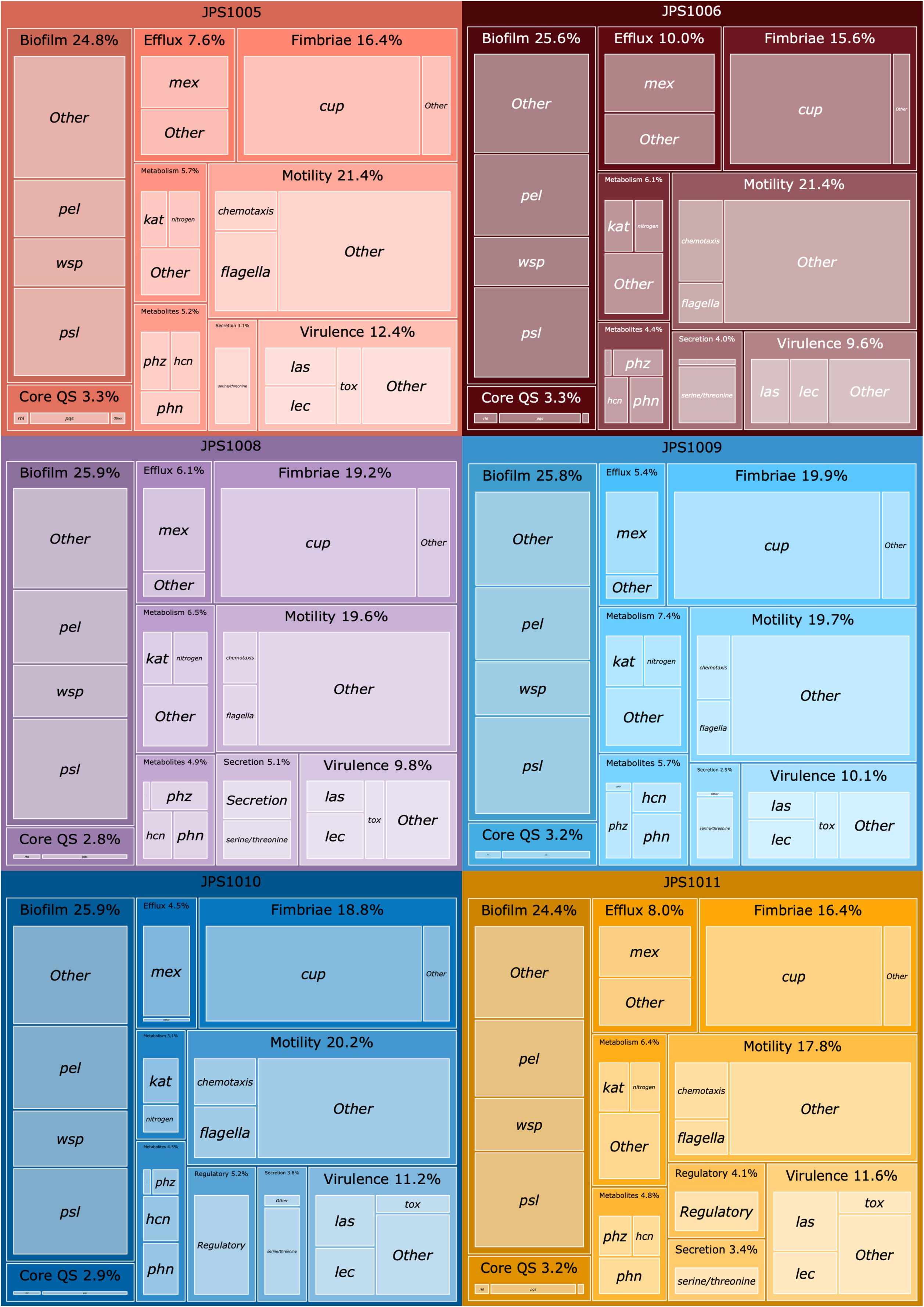
Categories of mutated genes and their function. The treemap was created using the list of called variants for each isolate filtered for non-synonymous mutations and 404 genes that are QS-regulated or QS-related. Percentage indicates the ratio of called variants in the functional category.

**Figure S2.**
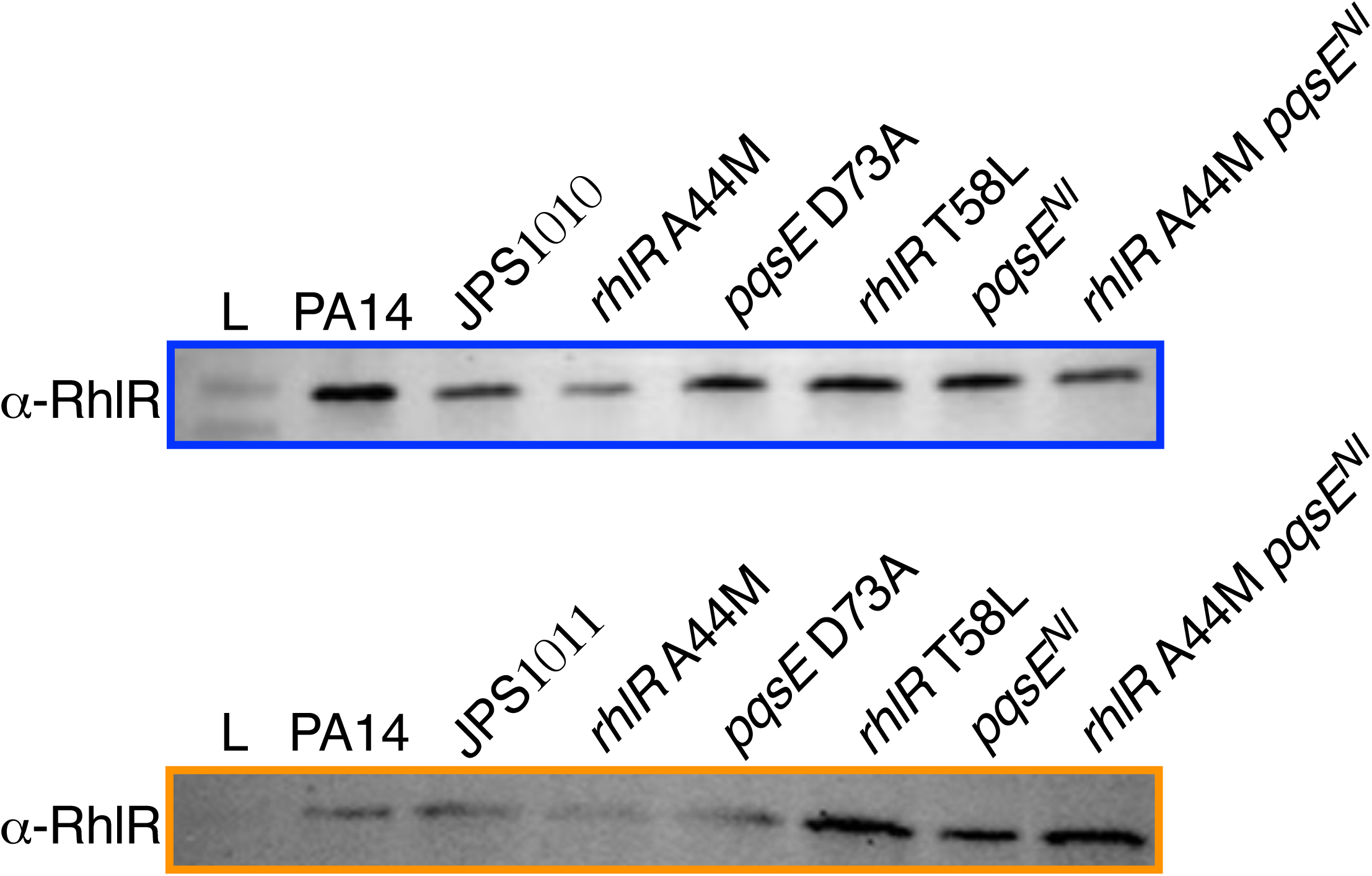
RhlR protein levels in clinical isolates and their mutant backgrounds. Western blot using a polyclonal antibody for RhlR against whole-cell lysates from JPS1010 (top), JPS1011 (bottom), and their respective isogenic mutants as indicated. A WT PA14 strain was used as a control.

## ACKNOWLEDGMENTS

The authors thank all members of the Paczkowski lab as well as the research laboratories in the Division of Genetics at the Wadsworth Center for helpful discussions on the research. The authors thank Dr. Alicia Mendoza (University at Albany) and Autumn Pope for critical reading of the manuscript as well as Dr. Lingyun Li (Wadsworth Center) for technical support during the mass spectrometry experiments. We thank Dr. Matthew Woflgang for the generous donation of isolates. This work was made possible with the help of the dedicated staff scientists at the Advanced Genomics Technologies Cluster and the Media & Tissue Core facilities at the Wadsworth Center. The authors acknowledge the contributions of Drs. Spencer Bruce and Erica Lasek-Nesselquist of the Bioinformatics Core and Dr. Jennifer Yates of the Immunology Core at the Wadsworth Center for training and technical support.

## FUNDING AND ADDITIONAL INFORMATION

This work was supported by National Institutes of Health training grant T32GM132066 to C.P.M., NIH grant R01GM14436101, New York Community Trust Foundation grant P19-000454, Cystic Fibrosis Foundation grant PACZKO21G0, and American Lung Association Innovation Award INALA2023 to J.E.P. The development of the cell culture model was supported by a SUNY Albany Grenander Award for Non-animal Methodologies in Research, Testing, and Education to M.L.S. and J.E.P. E.G.K., E.A.K., and L.M.K. are supported by Cooperative Agreement Number NU60OE000104 (CFDA #93.322), funded by the Centers for Disease Control and Prevention (CDC) of the US Department of Health and Human Services (HHS).

## ABBREVIATIONS AND NOMENCLATURE

AHL: acyl-homoserine lactone
AI: autoinducer
CF: cystic fibrosis
CFTR: cystic fibrosis transmembrane conductance receptor
LBP: ligand-binding pocket
pwCF: patient with cystic fibrosis
PqsE^NI^: PqsE non-interacting variant
SNP: single nucleotide polymorphism
QS: quorum sensing

